# Seasonal changes in fish eDNA signal vary between contrasting river types

**DOI:** 10.1101/2024.07.08.601838

**Authors:** Nathan P. Griffiths, Jonathan D. Bolland, Rosalind M. Wright, Petr Blabolil, James A. Macarthur, Graham S. Sellers, Bernd Hänfling

## Abstract

Due to the societal reliance on goods and services provided by river systems, and their close proximity to settlements, few modern-day rivers are without significant anthropogenic modifications. The natural river hydrology is often altered as a consequence of pumping water for flood alleviation, retaining water for irrigation and modifying channels for navigation. In recent years, water pumping stations have been found to have several adverse impacts, including fish mortality (direct and indirect) and habitat fragmentation. More broadly, modern-day river systems face a myriad of anthropogenic flow and channel modifications, with varying impacts on different fish life stages. To manage such risks in line with policy, knowledge of the overall fish community and priority species present is required. It is therefore important to understand the robustness of developing survey strategies across differently managed river systems. This study investigates the seasonal patterns of environmental DNA (eDNA) metabarcoding detections from water samples, taken across three differently managed river types over a one-year period. We observed some significant seasonal variation in detection rates and fish communities; however, this variation was not consistent among river types. Despite this, we found comparatively poor fish communities upstream of pumping stations all year-round, with pumped catchments containing significantly fewer species than the adjacent main river channel and our regional control site. Finally, we highlight that seasonal variation in detectability for the overall fish community may not always reflect that of priority species. In our case, we found favourable European eel (*Anguilla anguilla*) detection in the summer months across all river types. It is therefore recommended that rather than focusing on overall detectability, policy driven targeted surveys should be designed with priority species ecology in mind.

## 1. Introduction

European rivers are under broad scale pressures from hydromorphological alteration, resulting in habitat fragmentation and flow alteration (Szabolcs *et al*., 2022). This makes addressing and prioritising lotic freshwater conservation a challenge. Migratory species such as the European eel (*Anguilla anguilla*) are especially vulnerable to these pressures, and have faced significant declines over the last four decades (Aalto *et al*., 2016; Correia *et al*., 2018; ICES, 2019). In response, the EC Eel Regulation (1100/2007) was implemented, which requires the development of eel management plans (EMPs) aiming to achieve safe passage of >40% historic silver eel biomass between inland waters and the sea (Council of the European Union, 2007). However, with over one million barriers fragmenting European river systems (Belletti *et al*., 2020), it is important that mitigation measures can be prioritised to maximise the impacts of limited resources.

Due to the mounting evidence that water pumping stations can adversely impact fish communities in rivers (Norman *et al*., 2023a, 2023b; Solomon & Wright, 2012), particularly the critically endangered *A. anguilla* (Bolland *et al*., 2019; Buysse *et al*., 2014; Jacoby & Gollock, 2014), it is essential that we can prioritise management resources at these structures effectively. These impacts are direct and indirect, including fish mortality from blade strikes during downstream passage, and acting as barriers to both upstream and downstream migration (Bolland *et al*., 2019; Buysse *et al*., 2014, 2015; Kroes *et al*., 2020). However, in the absence of present-day knowledge of the fish community in such catchments, it is challenging to apply evidence-based prioritisation to mitigate these impacts (Solomon & Wright, 2012). Recent comparisons with environmental DNA (eDNA) metabarcoding have suggested that current standard practice surveys (seine netting / electrofishing) may under-represent low abundance priority species, including *A. anguilla,* in heavily managed catchments (Griffiths *et al*., 2020; McDevitt *et al*., 2019). Elsewhere, studies have also found that eDNA-based monitoring can provide a high sensitivity method of detecting priority species more generally (Burgoa Cardás *et al*., 2020; Halvorsen *et al*., 2020; McColl-Gausden *et al*., 2021; Muha *et al*., 2021; Weldon *et al*., 2020).

In order to implement eDNA-based monitoring into prioritisation frameworks in heavily managed river catchments, it is important that we first understand the dynamics of eDNA in such systems. It has recently been highlighted that in order to successfully integrate DNA-based methods into aquatic monitoring, best practice guidelines and standards must be developed (Blancher *et al*., 2022). Studies have found that eDNA metabarcoding can reflect long-term (Hänfling *et al*., 2016) and contemporary (Di Muri *et al*., 2020) fish catch data in lentic systems. Further, in lentic systems, eDNA signal has been found to be more homogenous in autumn/winter compared to the summer, and as a result detection probability is increased (Blabolil *et al*., 2022; Handley *et al*., 2019). Numerous studies have also found eDNA metabarcoding to yield high sensitivity in lotic systems (Griffiths *et al*., 2020; Hallam *et al*., 2021; McColl-Gausden *et al*., 2021; McDevitt *et al*., 2019; Muha *et al*., 2021; Pont *et al*., 2018), but detectability in rivers can be more unpredictable. Pumped catchments have artificial and highly variable flows (Kroes *et al*., 2020), switching from lentic (i.e. pumps off) to lotic (i.e. pumps on), thus making it difficult to predict how hydrology would influence eDNA assessments. It has been suggested that due to transportation of eDNA in rivers (Pont *et al*., 2018), high waterflow events can integrate more species and increase detection probability (Milhau *et al*., 2019). On the other hand, high flows have been observed to have conflicting effects, by diluting eDNA of rare species and reducing detectability (Curtis *et al*., 2021). Seasonal peaks in detectability of certain species have also been observed as a result of their ecology and lifecycle traits (i.e. spawning activity) (Bracken *et al*., 2019; Bylemans *et al*., 2017; Di Muri *et al*., 2022; Inui *et al*., 2021; Tillotson *et al*., 2018). Despite this, Milhau *et al* (2019) found that waterflow had a global influence on seasonal eDNA signal while reproductive period only influenced specific species. Ultimately, understanding seasonal patterns in lotic systems is more challenging, since temporal variations in hydrological conditions and species ecology can influence detectability, which may also vary depending on how flows are regulated and managed. As a result seasonal trends of eDNA-based monitoring have varied between different systems and target species (Burgoa Cardás *et al*., 2020; Curtis *et al*., 2021; Milhau *et al*., 2019; Sales *et al*., 2021; Wacker *et al*., 2019), and thus warrant further investigation in heavily managed catchments.

Given that the performance of eDNA-based monitoring is likely influenced by river type and target species ecology, it is important that such dynamics are considered before monitoring programmes are implemented. Here, we consider the seasonal patterns of detection using eDNA metabarcoding across three differently managed river types. This study aims to assess the comparability of eDNA metabarcoding seasonal detectability between different river types, provide insights into fish communities in small pumped catchments, and discuss the importance of targeted survey strategies. In order to determine this, we have the following objectives:

1. Assess the seasonal variation of eDNA metabarcoding detections over a one-year period across different river types.
2. Determine the impact of pumping stations on overall fish communities when compared to the main river channel and an unregulated regional control site.
3. Compare the seasonal detectability of priority species (*A. anguilla*) with that of the overall fish community.

## 2. Methods

### 2.1 Study systems

This study was carried out on the rivers Ancholme (Figure 1a) and Hull (Figure 1b), UK. Both flow into the Humber estuary, but differ considerably in geology, hydrology and management, and thus provide opportunity to evaluate the eDNA approach in contrasting lotic environments. The River Ancholme is typical of many lowland-streams in east-England, naturally slow flowing and heavily managed for navigation and drainage. The river level is controlled by a sluice gate at the river mouth, and there are 12 associated water pumping stations which are required to regulate water levels in tributaries / ditches which drain the surrounding land (i.e. pumped catchments). The River Hull is the most northern chalk stream in the UK and its flow is less intensively managed. The flow regime is typical of a chalk stream with relatively moderate spade conditions and moderated by heavy macrophyte cover. For the purpose of this study, we split our sampling plan to cover three distinct river types:

- River Ancholme pumped catchments (APC)
- Main River Ancholme (MRA)
- Main River Hull (MRH)

**Figure 1.**
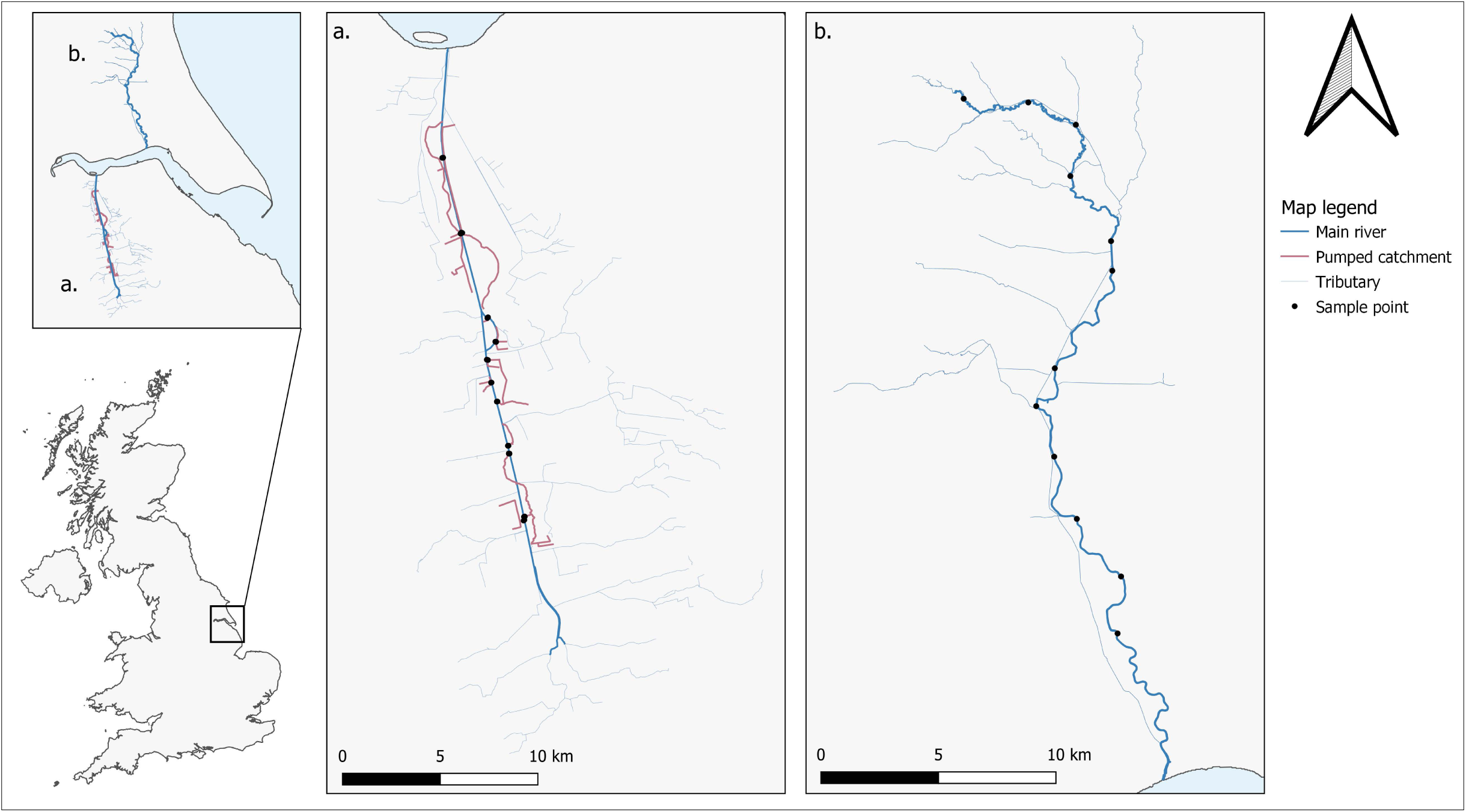
Map of the study sites, showing (a.) River Ancholme and (b.) River Hull catchments. eDNA sampling points are indicated by black dots (⦁). Note, River Ancholme samples were paired (upstream and downstream of pumping stations), therefore are represented by a single point on the map.

Twelve sample points were selected in each river type, allowing for four seasonal revisits over a one-year period. Sampling points were selected in the River Ancholme based on the position of the 12 water pumping stations present (Figure 1a), allowing paired upstream and downstream samples to be taken. In the River Hull, sampling points were allocated based on approximately equidistant access points (Figure 1b).

### 2.2 Water sampling

Across our study sites, 144 2L surface water samples were taken using sterile Gosselin HDPE plastic bottles (Fisher Scientific UK Ltd., Loughborough, UK). Each river type was sampled at twelve points (Figure 1) each season - based on the four UK meteorological seasons. This allowed for 48 samples to be taken from each river type, with the Ancholme and Hull catchments being sampled during Spring (11/03/2019, 27/05/2020), Summer (02/07/2019, 29/08/2019), Autumn (17/10/2019, 28/11/2019) and Winter (17/01/2020, 26/02/2020) respectively. Each 2L sample taken consisted of 5 x 400ml sub-samples spaced a few metres apart to account for any stochasticity in eDNA distribution within the watercourse. All samples were taken from the river bank, using a reach pole where required (Figure 2a). Sterile gloves were worn by the sampler and changed between samples, the sampling pole was cleaned with bleach (10%), and thoroughly rinsed between samples to prevent contamination. For each sampling visit, a 2L field blank (purified water) was taken out and handled alongside eDNA water samples to monitor for contamination.

**Figure 2.**
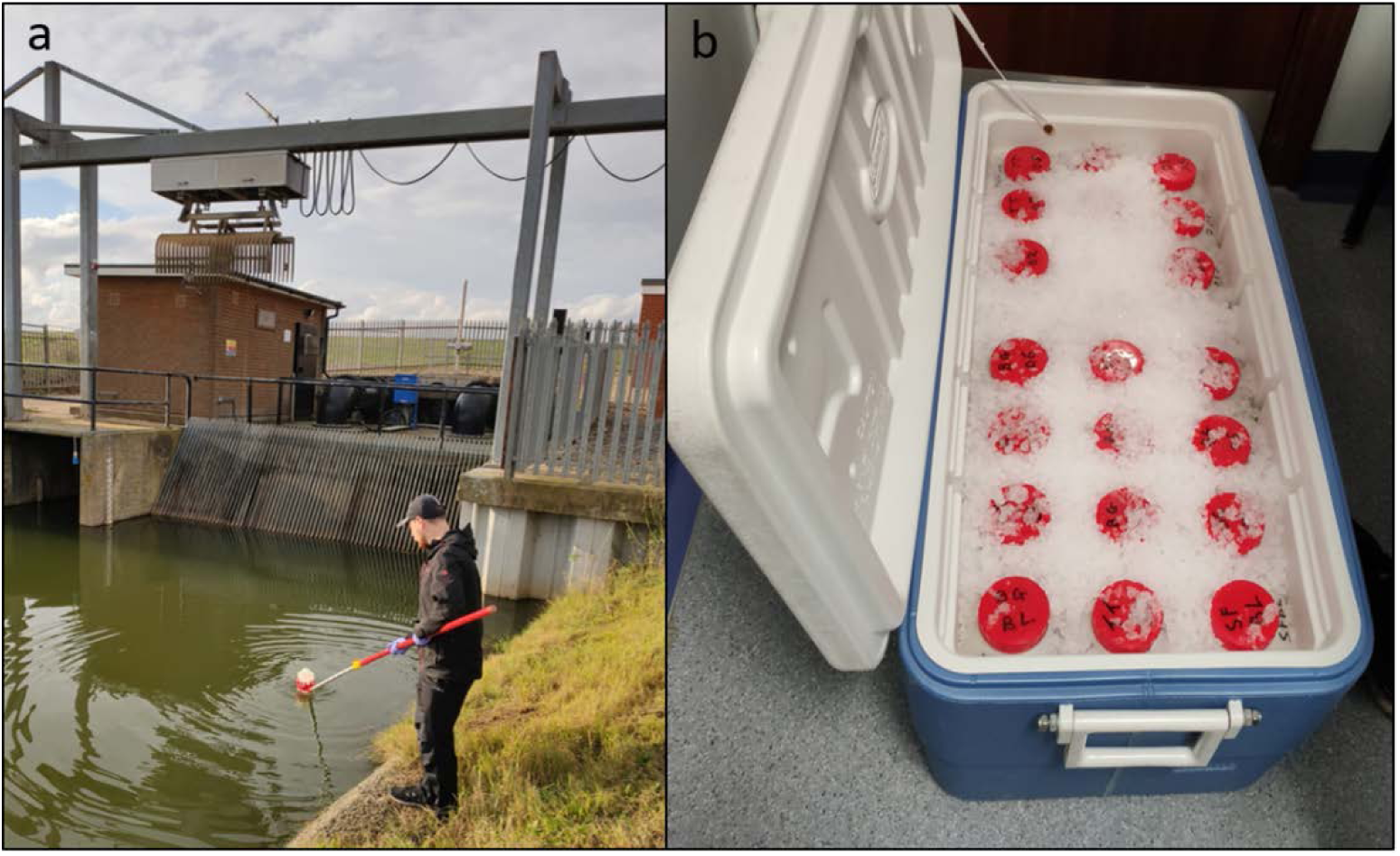
(a) Water sampling upstream of a pumping station using the reach pole and (b) water samples being stored on ice in a cool box until filtration.

Upon collection, water samples were immediately placed on ice in a bleach-sterilised cool box during transit and taken back to a dedicated eDNA filtration facility at the University of Hull (Figure 2b). All samples and blanks were vacuum filtered within 24h of the time collected. All surfaces and equipment in the filtration lab were sterilised using 10% v/v chlorine-based commercial bleach solution (Elliott Hygiene Ltd., Hull, UK). Between each filtration run, equipment was immersed in 10% bleach solution for 10 min, soaked in 5% v/v MicroSol detergent (Anachem, Leicester, UK) for an additional 10 min, and following this rinsed thoroughly with purified water to remove any detergent residue. When possible, the full 2L of water was vacuum filtered through sterile 0.45μm cellulose nitrate membrane filters with pads (47 mm diameter; Whatman, GE Healthcare, UK) using Nalgene filtration units – two filters were used per sample to reduce filter clogging. Once 1L of water had passed (or after 30 minutes under vacuum) the filters were removed from units using sterile tweezers, rolled, and placed back-to-back in sterile 5ml Axygen screw cap transport tubes (Fisher Scientific UK Ltd., Loughborough, UK) each pre-prepared with 1 g of 0.15 mm and 1 - 1.4 mm diameter sterile garnet beads ready for extraction (Sellers *et al*., 2018), then stored at −20°C.

### 2.3 DNA extraction

DNA from each sample was co-extracted from duplicate filters alongside extraction blanks at the University of Hull eDNA facility using a designated sterile extraction area, following the Mu-DNA: Water protocol (Sellers *et al*., 2018). Following extraction, the eluted DNA extracts (100μl) were quantified and checked for purity using a NanoDrop™ Spectrophotometer to confirm that DNA was isolated successfully, then stored at −20°C until PCR amplification.

### 2.4 Library preparation

The eDNA metabarcoding and library preparation workflow applied in this study follows that outlined in (Griffiths *et al*., 2023). This is summarised below:

Nested metabarcoding following a two-step PCR protocol was performed, using multiplex identification tags in both steps to enable sample identification as described in (Kitson *et al*., 2019). The first PCR (PCR1) was performed in triplicate (3× PCR replicates per eDNA extract), amplifying a 106bp fragment using published 12S ribosomal RNA primers 12S-V5-F (5′-ACTGGGATTAGATACCCC-3′) and 12S-V5-R (5′-TAGAACAGGCTCCTCTAG-3′) (Kelly *et al*., 2014; Riaz *et al*., 2011). These primers have been previously validated in silico, in vitro and in situ for UK freshwater fish, confirming that all UK species can be detected with the exceptions of distinctions between *Lampetra planeri*/*Lampetra fluviatilis* and *Perca fluviatilis*/*Sander lucioperca*; three species of Asian carp (*Hypophthalmichthys nobilis*, *Hypophthalmichthys molitrix* and *Ctenopharyngodon idella*); and species within the genera Salvelinus and Coregonus (Hänfling *et al*., 2016). PCR-negative controls (Molecular Grade Water) were used throughout, as were positive controls using DNA (0.05 ng μl^−1^) from the non-native cichlid *Maylandia zebra*. All PCR replicates were pooled, and samples from each PCR1 plate normalised and pooled to create sub-libraries. Sub-libraries were then purified using MagBIND RxnPure Plus magnetic beads (Omega Bio-tek Inc., Norcross, GA, USA), following a double size selection protocol (Quail *et al*., 2009). Ratios of 0.9× and 0.15× magnetic beads to 100μl of amplified DNA from each sub-library were used. Following this, a second shuttle PCR (PCR2) was performed on the cleaned product to bind Illumina adapters to the sub-libraries. A second purification was then carried out on the PCR2 products with Mag-BIND RxnPure Plus magnetic beads (Omega Bio-tek Inc., Norcross, GA, USA). Ratios of 0.7× and 0.15× magnetic beads to 50 μl of each sub-library were used. Eluted DNA was then refrigerated at 4°C until quantification and normalisation. Once pooled, the final library was then purified again (following the same protocol as the second clean-up), quantified by qPCR using the NEBNext Library Quant Kit for Illumina (New England Biolabs Inc., Ipswich, MA, USA) and verified for fragment size and purity using an Agilent 2200 TapeStation with High Sensitivity D1000 ScreenTape (Agilent Technologies, Santa Clara, CA, USA). Once verified, the library was loaded (mixed with 10% PhiX) and sequenced on an Illumina MiSeq using a MiSeq Reagent Kit v3 (600 cycle) (Illumina Inc., San Diego, CA, USA).

After sequencing, sub-libraries were demultiplexed to the sample level using a custom Python script. Tapirs, a reproducible workflow for the analysis of DNA metabarcoding data (https://github.com/EvoHull/Tapirs), was subsequently used for taxonomic assignment of demultiplexed reads. Sequence reads were quality trimmed, merged, and clustered before taxonomic assignment against a curated UK fish reference database (Hänfling *et al*., 2016). Note, that due to the minimum read length parameters selected for this study (90bp), Lampetra detections are not expected in the results. Taxonomic assignment here used a lowest common ancestor (LCA) approach based on basic local alignment search tool (BLAST) matches with minimum identity set at 98%.

### 2.5 Data processing and analysis

Data were analysed and visualised using R version 4.2.1 (R Core Team, 2020). A low-frequency reads threshold was applied to all eDNA samples prior to downstream analysis, which removed any reads making up less than 0.1% of total reads assigned to each sample as previously applied with this 12S marker (Handley *et al*., 2019; Hänfling *et al*., 2016). In addition to the low-frequency reads threshold, all assignments of *Cyprinus carpio* with fewer than 100 reads were omitted due to detections of this species in process blanks (3 - 65 reads). Samples with <500 reads were regarded as low quality and omitted from further analysis to reduce stochastic PCR effects. One site ‘ICAR’ from the APC was removed from analysis, due to instances of inhibition (samples <500 reads) and an artificial fishing pond directly upstream, meaning that detections were not representative of the fish community in the pumped catchments.

Prior to any statistical analysis, species richness data were screened for normality using base R. Differences in species richness between river types, and across seasons were tested for significance using ANOVA and nonparametric Kruskal-wallis tests, followed by post-hoc Tukey and Dunns tests respectively, depending on whether data conformed to normality. Differences in fish community between river types and across seasons were also tested using PERMANOVAs. Here, read counts were converted to presence/absence and Jaccard’s dissimilarity matrices computed. To enable comparison, outputs were visualised by Non-Metric multidimensional scaling (NMDS) and then differences between groupings tested for significance using PERMANOVA. Species accumulation curves including standard deviation based on 100 permutations were computed and visualised using the ‘specaccum’ function in the vegan package (Oksanen, 2013). Maps were created using QGIS software (QGIS Development Team, 2022). All additional plots and visualisations were made using ggplot2 in R (Wickham, 2016).

## 3. Results

### 3.1 Overall fish community between river types

When considering all samples taken across our study, the overall fish community varied between the three river types (Figure 3). The species present and site occupancy values were notably different between river types, with APC appearing to represent a diminished subset of the MRA, dominated by two stickleback species *Gasterosteus aculeatus* and *Pungitius pungitius* (Figure S2), while the overall composition of the MRH is visually different (Figure 3a). The total number of fish species detected across this study was 29 and there were significant differences in species richness between river types (X2 = 84.676, df = 2, p-value < 2.2e-16), as the APC (n = 14) supported fewer species than the MRA (n = 23) (post hoc Dunn test: Z = 8.29, P < 0.05) and MRH (n = 23) (Z = 7.74, P < 0.05) but MRA and MRH were similar (Z = 0.56, P = 0.58) (Figure 3c). Two species, sand goby (*Pomatoschistus minutus*) and topmouth gudgeon (*Pseudorasbora parva*) only appeared in a single sample across the whole study in MRH and APC, respectively, however were retained following the low-frequency reads threshold. Excluding single detections within each river type, the species richness retained for the APC = 9 (64.3%), MRA = 21 (91.3%), and MRH = 21 (91.3%). The overall fish community varied significantly between river types (PERMANOVA, R2 = 0.44, DF = 2, P = 0.001) (Figure 3d), and further inspection of the data suggest that this is driven by Nestedness in the APC (Figure S1).

**Figure 3.**
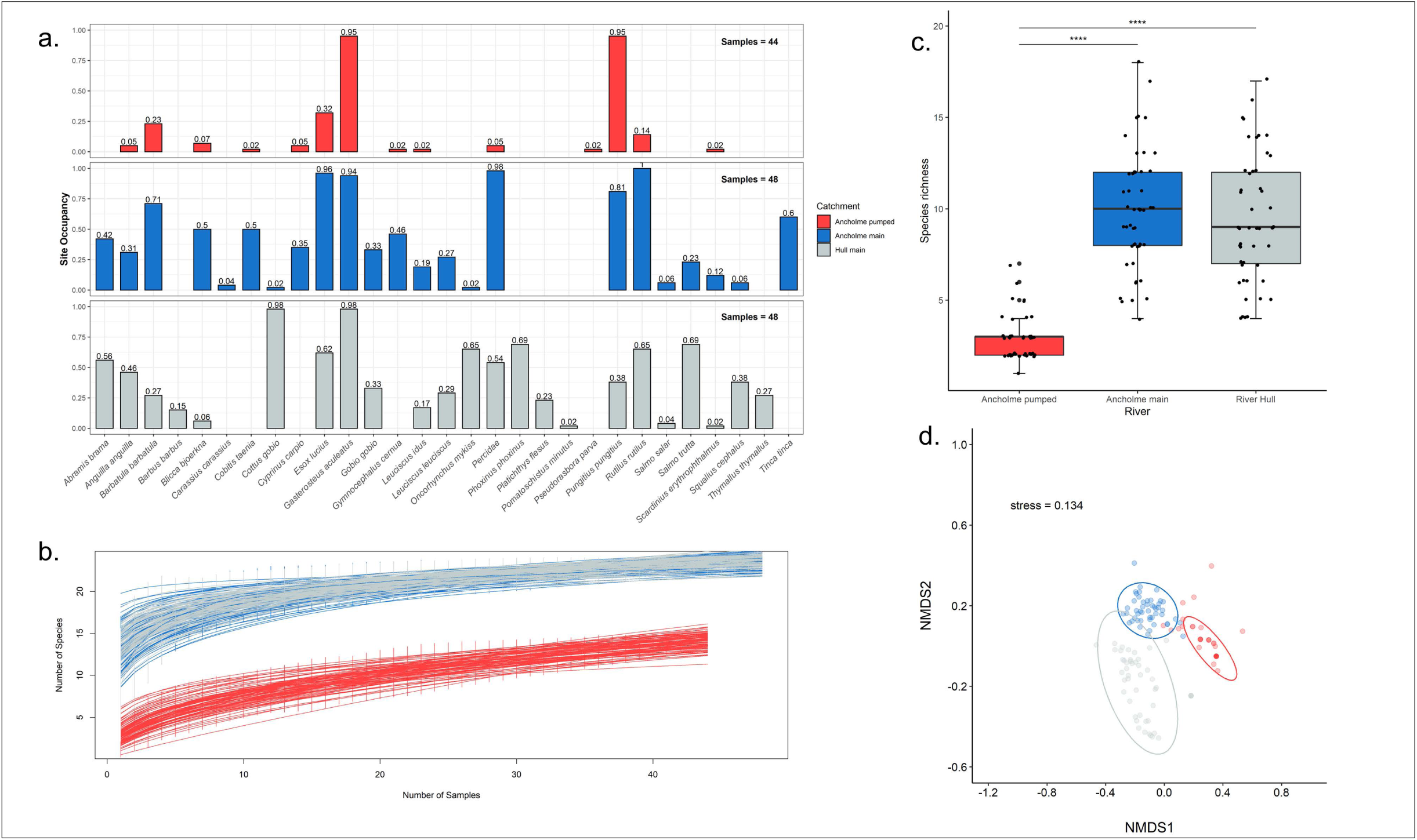
The overall fish community by river type, including the Ancholme pumped catchments (red), the main River Ancholme (blue) and the River Hull (grey). Plots visualise the following by river type; (a.) species site occupancy, (b.) species accumulation curves, (c.) species richness (**** = p<0.0001), (d.) NMDS of beta diversity based on Jaccard’s index including 95% confidence interval ellipses.

### 3.2 Seasonal dynamics of eDNA

#### 3.2.1 Ancholme pumped catchments (APC)

In the APC, 5/14 species (35.7%) were detected consistently across the four seasonal sampling events (Figure 4a). While no single sampling event detected all species, the highest species richness (n = 10) was in summer and the lowest (n = 7) in spring (Figure 4b). Overall, however, species richness was generally low and statistically comparable between seasons (X2 = 3.09, df = 3, p-value = 0.3781) (Figure 4c). The overall fish community also did not vary between seasons (PERMANOVA, R2 = 0.03, DF = 3, P = 0.982) (Figure 4d).

**Figure 4.**
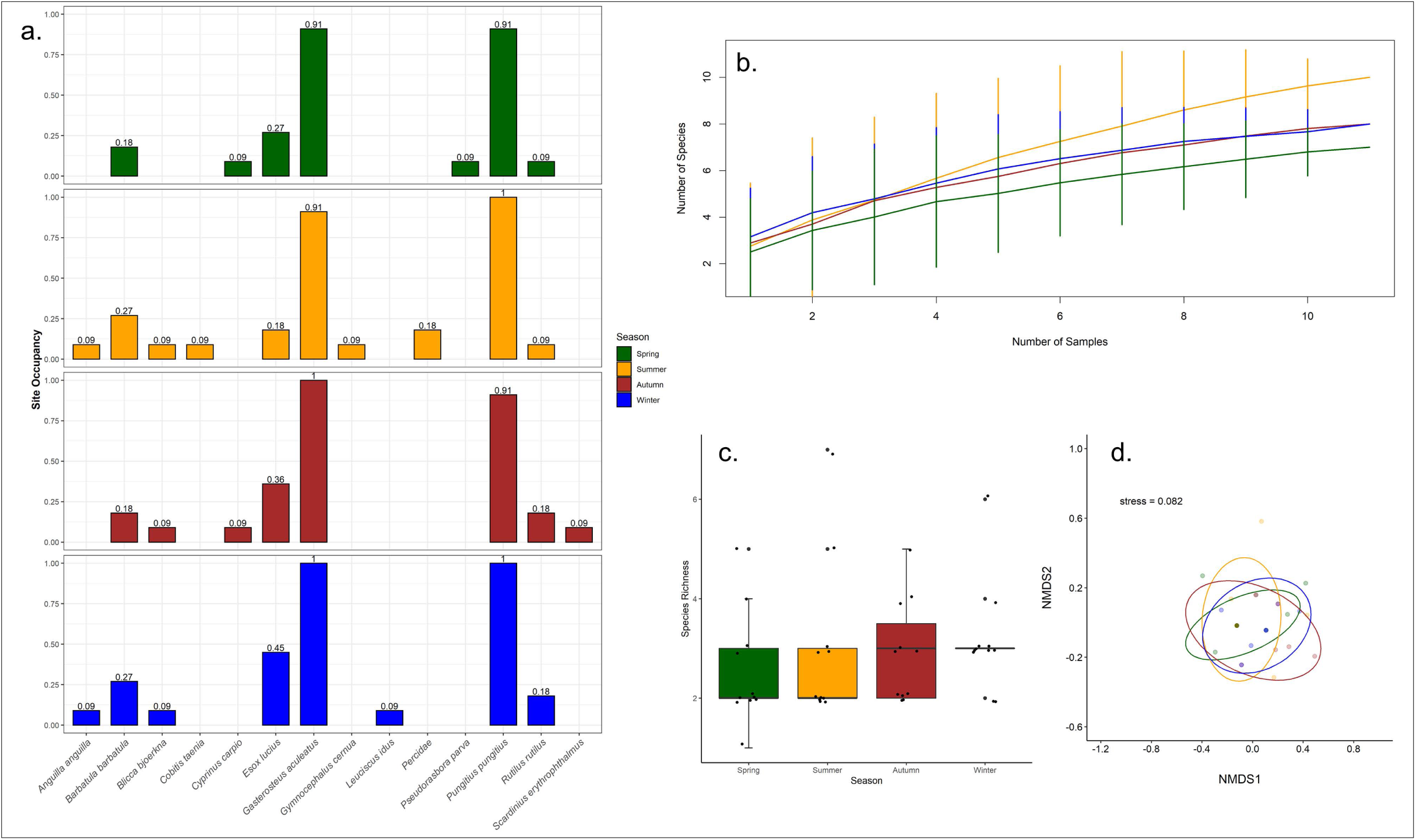
Fish community by season in the River Ancholme pumped catchment, including Spring (green), Summer (orange), Autumn (brown) and Winter (blue). Plots visualise the following by season; (a.) species site occupancy, (b.) species accumulation curves, (c.) species richness, (d.) NMDS of beta diversity based on Jaccard’s index including 95% confidence interval ellipses.

#### 3.2.2 Main River Ancholme (MRA)

In the MRA, 14/23 species (60.9%) were detected consistently across the four seasonal sampling events (Figure 5a). All 23 species were detected in winter while summer (n = 15) sampling detected the lowest overall species richness (Figure 5b). Species richness (ANOVA; F = 20.354, DF = 3, P < 0.0001) (Figure 5c) and overall fish community (PERMANOVA, R2 = 0.23, DF = 3, P = 0.001) (Figure 5d) varied significantly between seasons.

**Figure 5.**
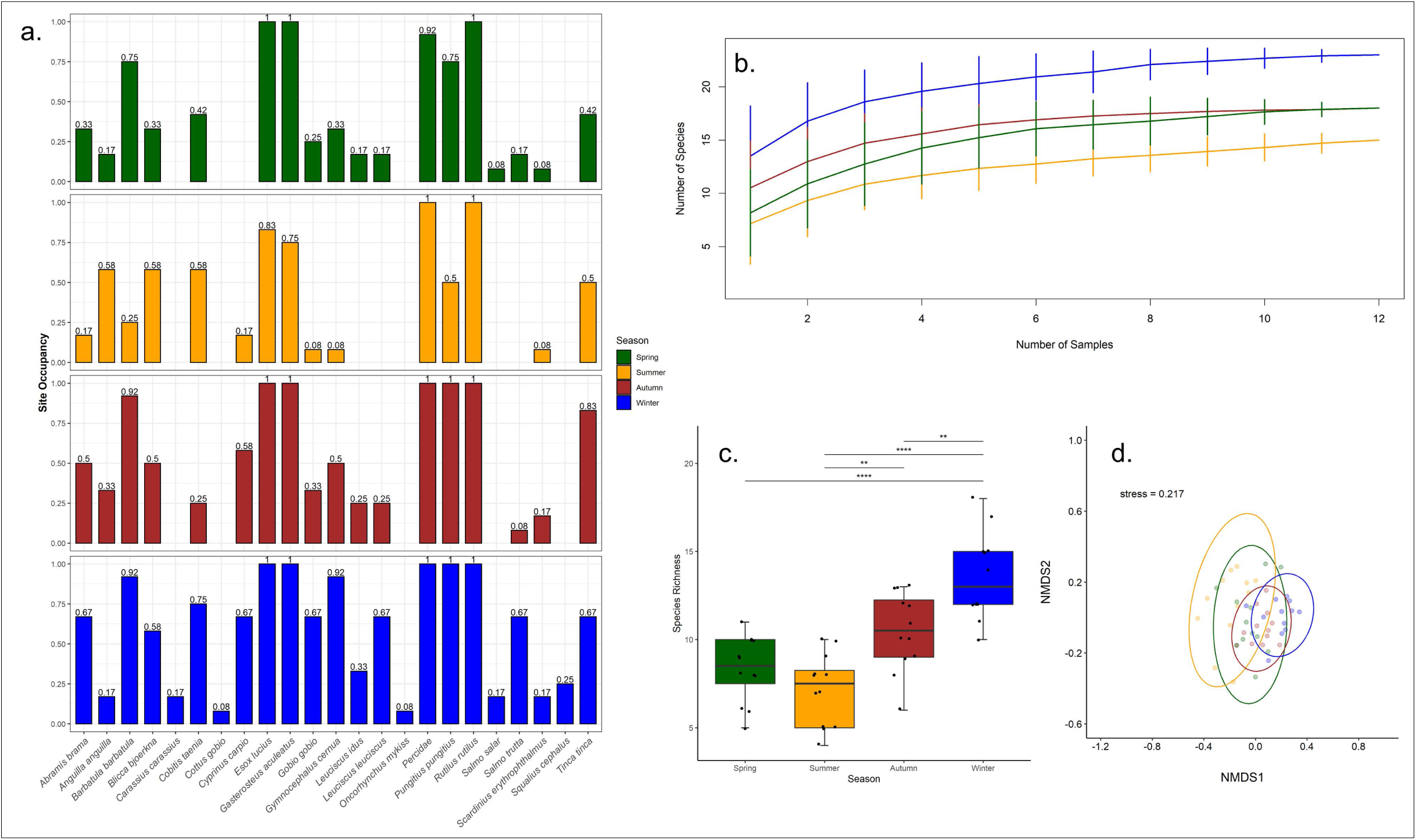
Fish community by season in the main River Ancholme, including Spring (green), Summer (orange), Autumn (brown) and Winter (blue). Plots visualise the following by season; (a.) species site occupancy, (b.) species accumulation curves, (c.) species richness (** = p<0.01, **** = p<0.0001), (d.) NMDS of beta diversity based on Jaccard’s index including 95% confidence interval ellipses.

#### 3.2.3 Main River Hull (MRH)

In the MRH, 16/23 species (69.6%) were detected consistently across the 4 sampling events (Figure 6a). While no single sampling event detected all species, the overall species richness was highest in summer (n = 21) and lowest in autumn (n = 16) (Figure 6b), but seasonal differences were not statistically significant (ANOVA; F = 2.3305, DF = 3, P = 0.0873) (Figure 6c). However, the overall fish community detected varied significantly between seasons (PERMANOVA, R2 = 0.12701, DF = 3, P = 0.015) (Figure 6d).

**Figure 6.**
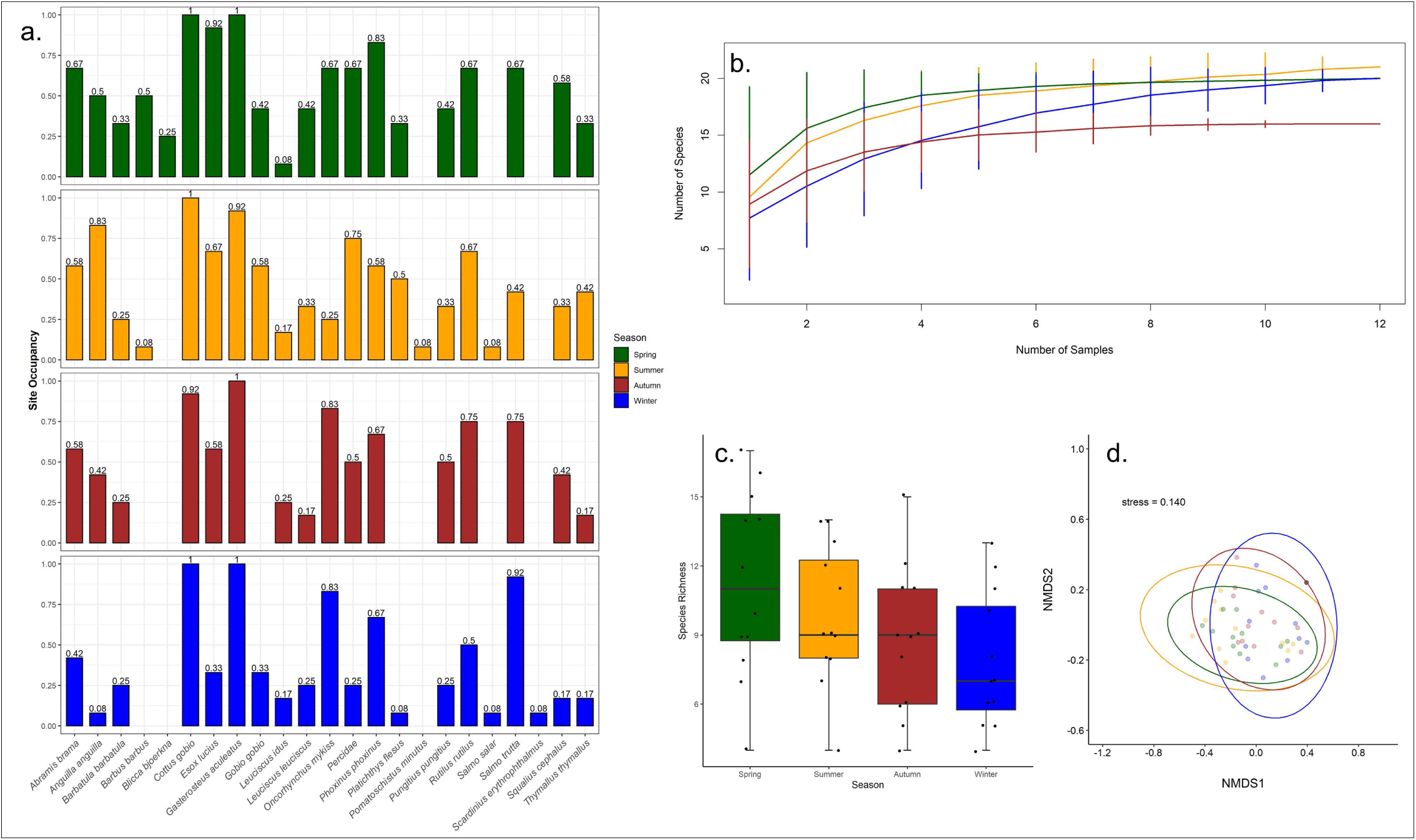
Fish community by season in the River Hull, including Spring (green), Summer (orange), Autumn (brown) and Winter (blue). Plots visualise the following by season; (a.) species site occupancy, (b.) species accumulation curves, (c.) species richness, (d.) NMDS of beta diversity based on Jaccard’s index including 95% confidence interval ellipses.

### 3.3 Priority species (*Anguilla anguilla*)

*A. anguilla* detections were consistently highest in summer between river types (Figure 7), i.e. 46.4% of eel detections in the MRA and 45.4% in the MRH. Consequently, across our study almost half (46.2%) of all eel detections (18/39) occurred in summer sampling events, which was significantly higher than 9/39 (23.1%) in autumn (p<0.05), 8/39 (20.5%) in spring (p<0.05), and 4/39 (10.3%) in winter (p<0.001) (Figure 7). Eels were only detected in two samples within the APC, both detections occurred at the same sampling site in summer and winter. In the MRH, average eel reads were significantly higher (p<0.001) in summer than winter sampling events (Figure 7).

**Figure 7.**
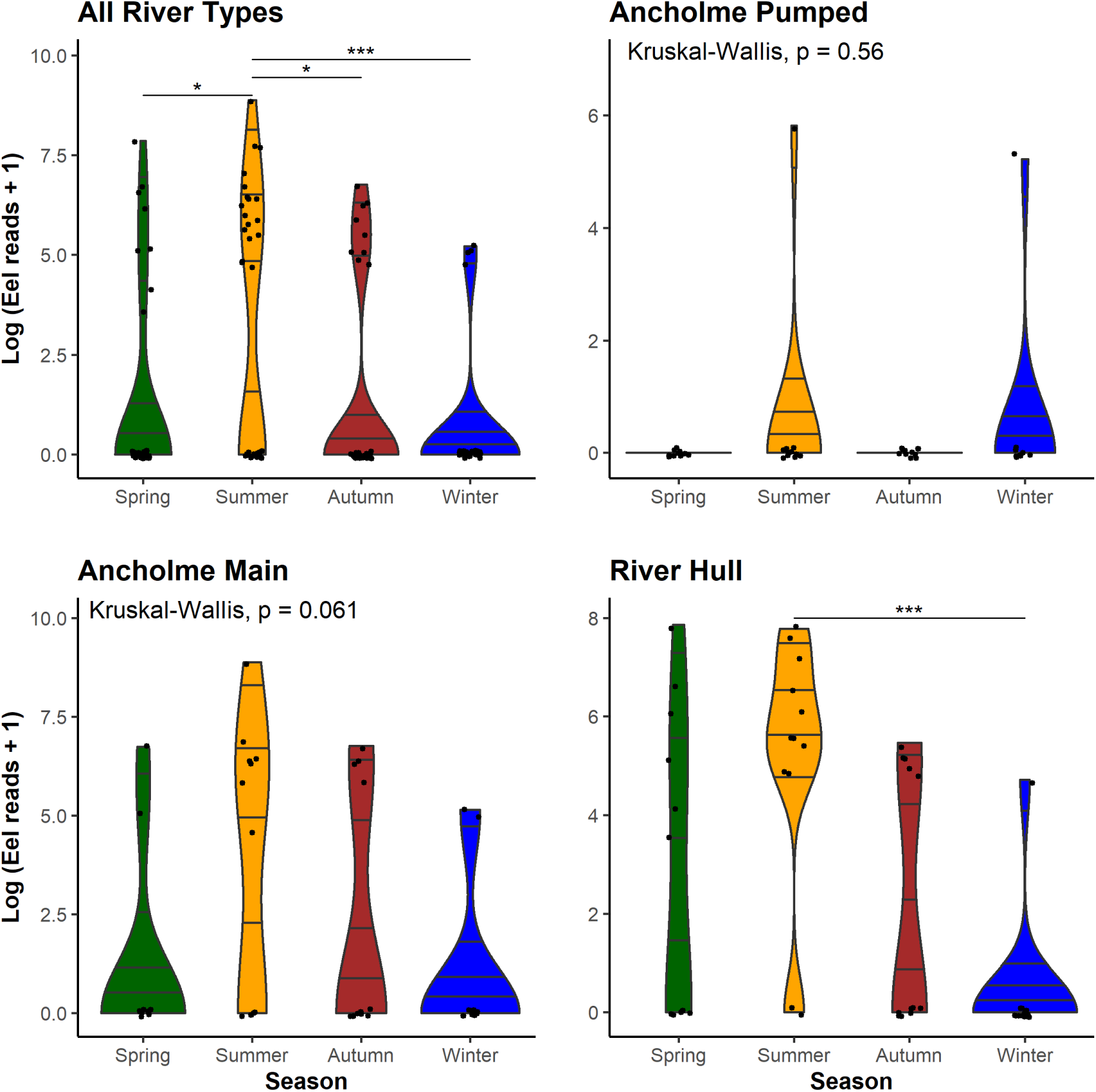
Violin plots visualising the differences in *A. anguilla* reads detected between seasons in all river types, and within each individual river type. Kruskal-Wallis tests were carried out to determine significant differences and displayed here, post hoc Dunn tests were carried out on significant outputs to determine the seasonal drivers of significance (* = p<0.05, *** = p<0.001). *Note – number of Log eDNA reads are not directly associated with eel abundance.

## 4. Discussion

This study for the first time compared the performance of fish eDNA metabarcoding among heavily managed and contrasting lotic environments; 12 tributaries with water pumping stations to regulate water levels (APC), a lowland river regulated by a sluice gate (MRA) and a chalk stream with man-made weirs (MRH). Variation in the seasonal performance of eDNA monitoring was assessed through comparing alpha diversity (i.e. average species richness per sampling occasion) and community similarity (i.e. Jaccard’s index). The three river types investigated in this study showed contrasting patterns of seasonal variation in eDNA-based fish detection. Overall species richness was not significantly different among seasons in the APC and MRH, but differed significantly in the MRA. Based on community similarity analysis there was no overall significant difference among samples from different seasons in the APC, but significant differences were observed in MRA and MRH. There was less seasonal variation in the MRH but marked seasonal differences in the MRA, while the APC showed a significantly impoverished fish fauna irrespective of the monitoring season. We also observed favourable detections of our priority species *A. anguilla* in summer across all river types, despite significantly higher overall species detectability in the MRA winter samples.

### 4.1 Seasonal dynamics of eDNA

The different seasonal responses in eDNA detectability among river types could be driven by differences in their fish communities or abiotic environments, especially hydrological conditions. Previous studies have shown seasonal peaks in detectability of certain species and that these are in turn influenced by their specific ecology and lifecycle traits, especially the timing of spawning activities (Bracken *et al*., 2019; Bylemans *et al*., 2017; Di Muri *et al*., 2022; Inui *et al*., 2021; Tillotson *et al*., 2018). This is one potential explanation for the patterns observed in our study, as fish communities varied significantly between the three river types (Figure 3), although the species present in APC were a subset of those detected in MRA (Figure S1). Additionally, the significantly higher detection rates observed in MRA winter samples appear to be driven by a broad increase in eDNA signal (Figure 5). This suggests that in this instance abiotic factors, including hydrological conditions, may be driving seasonal variation in detectability. Indeed, a study by (Milhau *et al*., 2019) found that waterflow had a global influence on eDNA-based taxonomic richness, while fish reproductive periods only influenced specific species. Further, it was suggested that sampling in periods of higher flow may be beneficial to obtain a more integrative overview of the fish community (Milhau *et al*., 2019). In addition, environmental DNA metabarcoding has been found to have increased detection efficiency in the autumn/winter for fish communities in mountain streams (Suzuki *et al*., 2022). Our data support this, with significantly increased detectability in winter samples in the MRA. Other studies however, have reported that increased flow can dilute eDNA concentrations, and therefore reduce detectability (Curtis *et al*., 2021).

More species were detected consistently across all seasons in MRH (69.6%) and MRA (60.9%) than APC (35.7%). Although, despite the poor consistency of low-frequency species detections in APC, there were no significant differences in species richness or fish community between sampling seasons (Figure 4). By contrast, in the more species rich MRA and MRH, the fish communities detected varied significantly between sampling seasons (Figure 5, Figure 6). While both were significant, the MRA community composition was much more variable than that of the MRH (Figure 5d, Figure 6d). Given that the MRA is heavily managed and channelised with a sluice gate regulating river levels at the mouth, flows are more variable. While the MRH is less intensively managed and is a chalk stream with more stable flows, it is still subject to seasonal variation in flow rate (supplementary information). It is possible that the significant differences in seasonal detection in MRA are also driven by these increased fluctuations in hydrological conditions, however further research into this is required. In addition, water temperature, UV light and pH levels can all influence the shedding and degradation rates of eDNA (Buxton *et al*., 2017; Strickler *et al*., 2015). This could lead to more patchy distributions of eDNA, in the absence of high flows leading to mixing and downstream transportation (Pont *et al*., 2018). With this in mind, it is likely that seasonal changes in fish community detections are influenced by many interacting factors, which may be amplified in highly variable catchments.

### 4.2 Fish community in pumped catchments

Fish communities varied significantly between the three river types with significantly less species detected in the APC compared to MRA and our regional control site MRH (Figure 3), although the species present in APC were a subset of those detected in MRA (Figure S1). Furthermore, one consistent characteristic of APC was the low diversity fish community dominated by the stickleback species *G. aculeatus* and *P. pungitius* which indicates poor habitat quality. It is already well established that pumping stations can lead to fish mortality during downstream passage (Buysse *et al*., 2014, 2015; Kroes *et al*., 2020)ore recently, (Norman *et al*., 2023b) found extreme flood-relief pump operations significantly altered resident fish populations due to heavily degraded longitudinal habitat and isolated lateral connectivity limiting access to flow refuge. In addition, pumping stations often act as barriers to upstream fish movement, thereby hindering lateral connectivity with the main river (Bolland *et al*., 2012; Manfrin *et al*., 2020), including re-colonisation following extreme pumping events.

### 4.3 Targeting priority species (*Anguilla anguilla*)

Despite the variation in overall detectability between river types, we found that *A. anguilla* detection was consistently favourable in summer and lowest in winter in MRA and MRH (Figure 7). This is in contrast with a recent study by (Burgoa Cardás *et al*., 2020), which found increased eel detections associated with their upstream migration period (February – April) and reduced detection in the summer months (July). The aforementioned study however attributed these increased detection rates to glass eel recruitment leading to additional positive sites downstream in the system, while upstream resident eels were detected consistently through the seasons. In our study, peaks in summer could be due to increased activity and DNA shedding rate of resident eels in conjunction with slower flows allowing eDNA of rare/low abundance species to accumulate by reducing the dilution effect associated with high flows (Curtis *et al*., 2021). Additionally, resident eels display overwintering dormancy behaviour in colder temperatures, where they avoid shallow areas and remain motionless (Westerberg & Sjöberg, 2015), likely reducing eDNA shedding. These contrasting results are an example of how eDNA dynamics may not always be uniform across different workflows, river systems and target species. Regarding the APC, only two detections of *A. anguilla* occurred across our study, both upstream of the same pumping station. This could suggest an element of increased connectivity at this structure given the catadromous life cycle of *A. anguilla*, and thus, its absence upstream of 11/12 pumping stations further highlights the lack of upstream passability. This finding is an example of how eDNA-based methods can be applied to prioritise structures for eel regulations for example. In addition, we would recommend summer months as the optimal sampling period for *A. anguilla* in such modified river systems.

## 5. Conclusions

Our findings support that seasonal variation in eDNA patterns are not always comparable between lotic systems, and that this should be taken into consideration when designing and planning future studies and monitoring programmes. In addition, flow management is likely to be a contributing factor in this, since it has an impact on eDNA transport and concentration. Previous studies have found conflicting results between seasonal eDNA signals, and our metabarcoding workflow responded differently in our three river types. This has been observed in species specific analysis, where (Curtis *et al*., 2021) highlighted that such factors may be as or more important than the density of target species when explaining eDNA outputs. It is acknowledged that eDNA production, persistence and movement will differ by system and target species (Klymus *et al*., 2015), and other studies have observed seasonal changes in fish assemblages using eDNA metabarcoding (Milhau *et al*., 2019; Sales *et al*., 2021; Suzuki *et al*., 2022). Our findings highlight that seasonal variation in eDNA performance for detecting fish communities is not always transferable between contrasting river systems. We also highlight the impact of pumping stations in heavily managed river catchments, leading to an impoverished fish community in comparison with the associated main river channel. While overall seasonal detectability did vary between river types, detection of our target species *A. anguilla* was favourable in summer, and significantly higher than winter in our naturally flowing control site. Based on these results, we recommend that both biotic (i.e. specific species ecology) and abiotic (i.e. flow modifications) factors of heavily managed river systems should be considered when designing sampling strategies, particularly surrounding the anthropogenic modifications to hydrodynamics and spatial resolution requirements for sampling. Given the highly modified nature of European river systems (Belletti *et al*., 2020; Szabolcs *et al*., 2022), and increasing drive for eDNA integration into aquatic monitoring (Blancher *et al*., 2022), we emphasise that future research regarding the impact of specific modification regimes on eDNA dynamics should be considered to aid interpreting results for management purposes.

## DATA AVAILABILITY STATEMENT

All scripts and corresponding data have been archived and made available at Zenodo: https://doi.org/10.5281/zenodo.12629881

## References

Aalto, E., Capoccioni, F., Terradez Mas, J., Schiavina, M., Leone, C., De Leo, G., & Ciccotti, E. (2016). Quantifying 60 years of declining European eel (Anguilla anguilla L., 1758) fishery yields in Mediterranean coastal lagoons. ICES journal of marine science: journal du conseil, 73, 101–110.

Belletti, B., Garcia de Leaniz, C., Jones, J., Bizzi, S., Börger, L., Segura, G.,… Zalewski, M. (2020). More than one million barriers fragment Europe’s rivers. Nature, 588, 436–441.

Blabolil, Griffiths, Hänfling, Jůza, Draštík, Knežević-Jarić,… Peterka. (2022). The true picture of environmental DNA, a case study in harvested fishponds. Ecological indicators, 142, 109241.

Blancher, P., Lefrançois, E., Rimet, F., Vasselon, V., Argillier, C., Arle, J.,… Bouchez, A. (2022). A strategy for successful integration of DNA-based methods in aquatic monitoring. Metabarcoding and Metagenomics, 6, e85652.

Bolland, J. D., Nunn, A. D., Lucas, M. C., & Cowx, I. G. (2012). The importance of variable lateral connectivity between artificial floodplain waterbodies and river channels: Rehabilitation of floodplain connectivity. River research and applications, 28, 1189– 1199.

Bolland, J. D., Murphy, L. A., Stanford, R. J., Angelopoulos, N. V., Baker, N. J., Wright, R. M.,… Cowx, I. G. (2019). Direct and indirect impacts of pumping station operation on downstream migration of critically endangered European eel. Fisheries management and ecology, 26, 76–85.

Bracken, F. S. A., Rooney, S. M., Kelly-Quinn, M., King, J. J., & Carlsson, J. (2019). Identifying spawning sites and other critical habitat in lotic systems using eDNA ‘snapshots’: A case study using the sea lamprey Petromyzon marinus L. Ecology and evolution, 9, 553–567.

Burgoa Cardás, J., Deconinck, D., Márquez, I., Peón Torre, P., Garcia-Vazquez, E., & Machado-Schiaffino, G. (2020). New eDNA based tool applied to the specific detection and monitoring of the endangered European eel. Biological conservation, 250, 108750.

Buxton, A. S., Groombridge, J. J., Zakaria, N. B., & Griffiths, R. A. (2017). Seasonal variation in environmental DNA in relation to population size and environmental factors. Scientific reports, 7, 46294.

Buysse, D., Mouton, A. M., Stevens, M., Van den Neucker, T., & Coeck, J. (2014). Mortality of European eel after downstream migration through two types of pumping stations. Fisheries management and ecology, 21, 13–21.

Buysse, D., Mouton, A. M., Baeyens, R., & Coeck, J. (2015). Evaluation of downstream migration mitigation actions for eel at an Archimedes screw pump pumping station. Fisheries management and ecology, 22, 286–294.

Bylemans, J., Furlan, E. M., Hardy, C. M., McGuffie, P., Lintermans, M., & Gleeson, D. M. (2017). An environmental DNA-based method for monitoring spawning activity: a case study, using the endangered Macquarie perch (Macquaria australasica). Methods in ecology and evolution / British Ecological Society, 8, 646–655.

Correia, M. J., Costa, J. L., Antunes, C., De Leo, G., & Domingos, I. (2018). The decline in recruitment of the European eel: new insights from a 40-year-long time-series in the Minho estuary (Portugal). ICES journal of marine science: journal du conseil.

Council of the European Union. (2007). Council Regulation (EC) No 1100/2007 of 18 September 2007 establishing measures for the recovery of the stock of European eel. Official Journal of the European Union L, 248.

Curtis, A. N., Tiemann, J. S., Douglass, S. A., Davis, M. A., & Larson, E. R. (2021). High stream flows dilute environmental DNA (eDNA) concentrations and reduce detectability. Diversity & distributions, 27, 1918–1931.

Di Muri, C., Lawson Handley, L., Bean, C. W., Li, J., Peirson, G., Sellers, G. S.,… Hänfling, B. (2020). Read counts from environmental DNA (eDNA) metabarcoding reflect fish abundance and biomass in drained ponds. Metabarcoding and Metagenomics, 4, e56959.

Di Muri, C., Lawson Handley, L., Bean, C. W., Benucci, M., Harper, L. R., James, B.,… Hänfling, B. (2022). Spatio-temporal monitoring of lake fish spawning activity using environmental DNA metabarcoding. Environmental DNA, n/a.

Griffiths, N. P., Bolland, J. D., Wright, R. M., Murphy, L. A., Donnelly, R. K., Watson, H. V., & Hänfling, B. (2020). Environmental DNA metabarcoding provides enhanced detection of the European eel Anguilla anguilla and fish community structure in pumped river catchments. Journal of fish biology, 97, 1375–1384.

Griffiths, N. P., Wright, R. M., & Hänfling, B. (2023). Integrating environmental DNA monitoring to inform eel (Anguilla anguilla) status in freshwaters at their easternmost range—A case study in Cyprus. Ecology.

Hallam, J., Clare, E. L., Jones, J. I., & Day, J. J. (2021). Biodiversity assessment across a dynamic riverine system: A comparison of eDNA metabarcoding versus traditional fish surveying methods. Environmental DNA, 3, 1247–1266.

Halvorsen, S., Korslund, L., Gustavsen, P. Ø., & Slettan, A. (2020). Environmental DNA analysis indicates that migration barriers are decreasing the occurrence of European eel (Anguilla anguilla) in distance from the sea. Global Ecology and Conservation, 24, e01245.

Handley, L. L., Read, D. S., Winfield, I. J., Kimbell, H., Johnson, H., Li, J.,… Hänfling, B. (2019). Temporal and spatial variation in distribution of fish environmental DNA in England’s largest lake. Environmental DNA. 2019, doi:10.1002/edn3.5.

Hänfling, B., Lawson Handley, L., Read, D. S., Hahn, C., Li, J., Nichols, P.,… Winfield, I. J. (2016). Environmental DNA metabarcoding of lake fish communities reflects long-term data from established survey methods. Molecular ecology, 25, 3101–3119.

ICES. (2019). Joint EIFAAC/ICES/GFCM Working Group on Eels (WGEEL). ICES Scientific Reports, 1, 177.

Inui, R., Akamatsu, Y., Kono, T., Saito, M., Miyazono, S., & Nakao, R. (2021). Spatiotemporal Changes of the Environmental DNA Concentrations of Amphidromous Fish Plecoglossus altivelis altivelis in the Spawning Grounds in the Takatsu River, Western Japan. Frontiers in Ecology and Evolution, 9.

Jacoby, D., & Gollock, M. (2014). Anguilla anguilla. The IUCN Red List of Threatened Species 2014: e. T60344A45833138. 2014.

Kelly, R. P., Port, J. A., Yamahara, K. M., & Crowder, L. B. (2014). Using environmental DNA to census marine fishes in a large mesocosm. PloS one, 9, e86175.

Kitson, J. J. N., Hahn, C., Sands, R. J., Straw, N. A., Evans, D. M., & Lunt, D. H. (2019). Detecting host–parasitoid interactions in an invasive Lepidopteran using nested tagging DNA metabarcoding. Molecular ecology, 28, 471–483.

Klymus, K. E., Richter, C. A., Chapman, D. C., & Paukert, C. (2015). Quantification of eDNA shedding rates from invasive bighead carp Hypophthalmichthys nobilis and silver carp Hypophthalmichthys molitrix. Biological conservation, 183, 77–84.

Kroes, R., Van Loon, E. E., Goverse, E., Schiphouwer, M. E., & Van der Geest, H. G. (2020). Attraction of migrating glass eel (Anguilla anguilla) by freshwater flows from water pumping stations in an urbanized delta system. The Science of the total environment, 714, 136818.

Manfrin, A., Bunzel-Drüke, M., Lorenz, A. W., Maire, A., Scharf, M., Zimball, O., & Stoll, S. (2020). The effect of lateral connectedness on the taxonomic and functional structure of fish communities in a lowland river floodplain. The Science of the total environment, 719, 137169.

McColl-Gausden, E. F., Weeks, A. R., Coleman, R. A., Robinson, K. L., Song, S., Raadik, T. A., & Tingley, R. (2021). Multispecies models reveal that eDNA metabarcoding is more sensitive than backpack electrofishing for conducting fish surveys in freshwater streams. Molecular ecology, 30, 3111–3126.

McDevitt, A. D., Sales, N. G., Browett, S. S., Sparnenn, A. O., Mariani, S., Wangensteen, O. S.,… Benvenuto, C. (2019). Environmental DNA metabarcoding as an effective and rapid tool for fish monitoring in canals. Journal of fish biology, 95, 679–682.

Milhau, T., Valentini, A., Poulet, N., Roset, N., Jean, P., Gaboriaud, C., & Dejean, T. (2019). Seasonal dynamics of riverine fish communities using eDNA. Journal of fish biology.

Muha, T. P., Rodriguez-Barreto, D., O’Rorke, R., Garcia de Leaniz, C., & Consuegra, S. (2021). Using eDNA Metabarcoding to Monitor Changes in Fish Community Composition After Barrier Removal. Frontiers in Ecology and Evolution, 9.

Norman, J., Reeds, J., Wright, R. M., & Bolland, J. D. (2023a). Impact of anthropogenic infrastructure on aquatic and avian predator–prey interactions in a modified lowland river. Freshwater biology.

Norman, J., Reeds, J., Wright, R. M., & Bolland, J. D. (2023b). The impact of extreme flood-relief pump operations on resident fish in an artificial drain and the potential for artificial habitat introduction. Fisheries management and ecology, 30, 483–493.

Oksanen, J. (2013). Vegan: ecological diversity. R Project.

Pont, D., Rocle, M., Valentini, A., Civade, R., Jean, P., Maire, A.,… Dejean, T. (2018). Environmental DNA reveals quantitative patterns of fish biodiversity in large rivers despite its downstream transportation. Scientific reports, 8, 10361.

QGIS Development Team. (2022). QGIS Geographic Information System. QGIS Association.

Quail, M. A., Swerdlow, H., & Turner, D. J. (2009). Improved protocols for the illumina genome analyzer sequencing system. Current protocols in human genetics / editorial board, Jonathan L. Haines… [et al.], Chapter 18, Unit 18.2.

R Core Team. (2020). R: A Language and Environment for Statistical Computing. Vienna, Austria: R Foundation for Statistical Computing 2020.

Riaz, T., Shehzad, W., Viari, A., Pompanon, F., Taberlet, P., & Coissac, E. (2011). ecoPrimers: inference of new DNA barcode markers from whole genome sequence analysis. Nucleic acids research, 39, e145.

Sales, N. G., Wangensteen, O. S., Carvalho, D. C., Deiner, K., Præbel, K., Coscia, I.,… Mariani, S. (2021). Space-time dynamics in monitoring neotropical fish communities using eDNA metabarcoding. The Science of the total environment, 754, 142096.

Sellers, G. S., Di Muri, C., Gómez, A., & Hänfling, B. (2018). Mu-DNA: a modular universal DNA extraction method adaptable for a wide range of sample types. Metabarcoding and Metagenomics, 2, e24556.

Solomon, J., & Wright, R. (2012). Prioritising pumping stations for facilities for the passage of eels and other fish. Environment Agency, Anglian Region.

Strickler, K. M., Fremier, A. K., & Goldberg, C. S. (2015). Quantifying effects of UV-B, temperature, and pH on eDNA degradation in aquatic microcosms. Biological conservation, 183, 85–92.

Suzuki, J., Nakano, D., & Kobayashi, S. (2022). Characteristics of diurnal and seasonal changes in fish detection patterns using environmental DNA metabarcoding in a mountain stream. Limnologica. 2022, p. 125955, doi:10.1016/j.limno.2022.125955.

Szabolcs, M., Kapusi, F., Carrizo, S., Markovic, D., Freyhof, J., Cid, N.,… Lengyel, S. (2022). Spatial priorities for freshwater biodiversity conservation in light of catchment protection and connectivity in Europe. PloS one, 17, e0267801.

Tillotson, M. D., Kelly, R. P., Duda, J. J., Hoy, M., Kralj, J., & Quinn, T. P. (2018). Concentrations of environmental DNA (eDNA) reflect spawning salmon abundance at fine spatial and temporal scales. Biological conservation, 220, 1–11.

Wacker, S., Fossøy, F., Larsen, B. M., Brandsegg, H., Sivertsgård, R., & Karlsson, S. (2019). Downstream transport and seasonal variation in freshwater pearl mussel (Margaritifera margaritifera) eDNA concentration. Environmental DNA, 1, 64–73.

Weldon, L., O’Leary, C., Steer, M., Newton, L., Macdonald, H., & Sargeant, S. L. (2020). A comparison of European eel Anguilla anguilla eDNA concentrations to fyke net catches in five Irish lakes. Environmental DNA, 57, 109.

Westerberg, H., & Sjöberg, N. (2015). Overwintering dormancy behaviour of the European eel (Anguilla anguillaL.) in a large lake. Ecology of freshwater fish, 24, 532–543.

Wickham, H. (2016). ggplot2: Elegant Graphics for Data Analysis. Springer-Verlag New York.

