## supplementary information for "Seasonal changes in fish eDNA signal vary between contrasting river types"

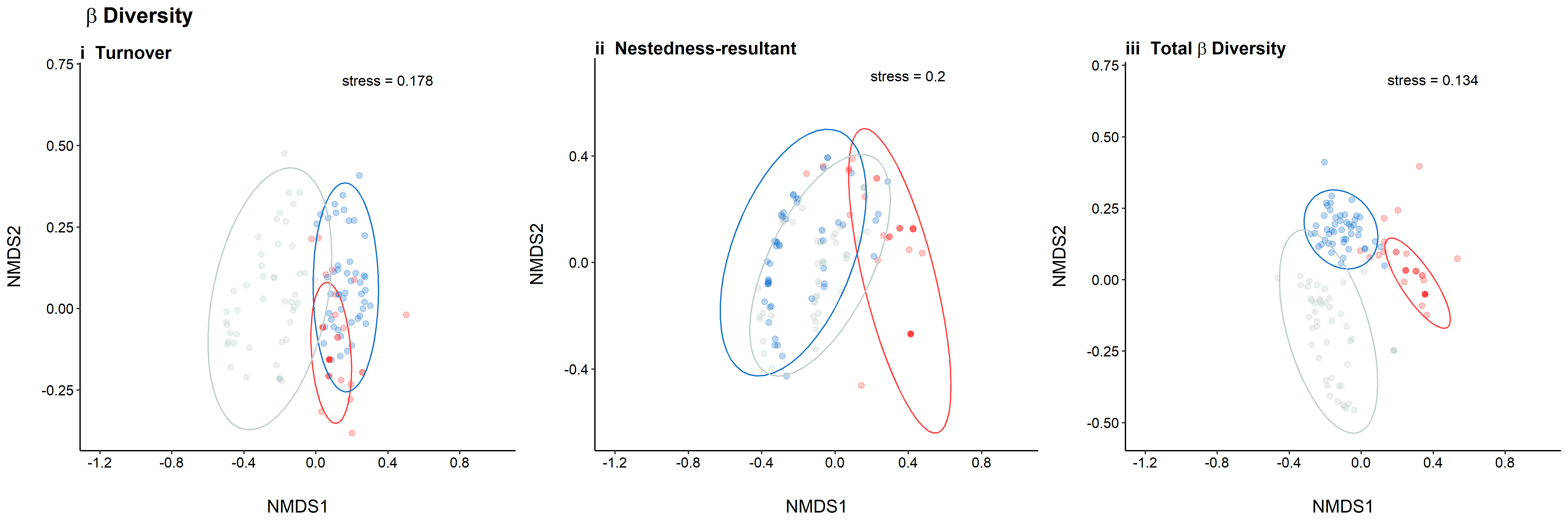

**Figure S1.** NMDS representing the driving components and differences in beta diversity between river types, Ancholme Pumped (red), Main Ancholme (blue), and River Hull (grey). From left to right, these plots represent differences in Turnover, Nestedness, and total beta diversity including 95% confidence interval ellipses.

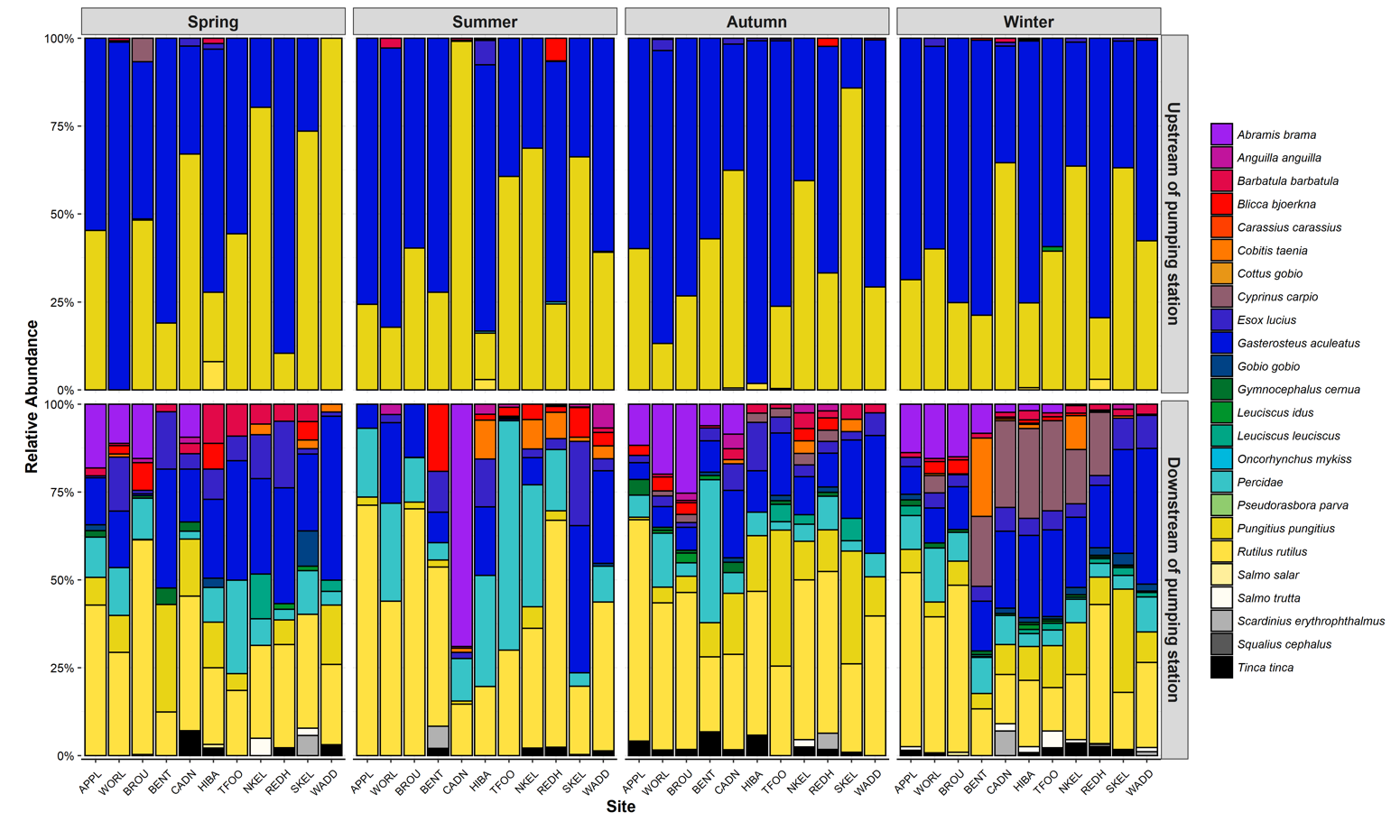

**Figure S2.** Bar plots comparing river types in the Ancholme catchment, including Ancholme Pumped catchments (top) and Ancholme main (bottom). Species composition is shown for each site represented by relative abundance (% reads), and repeat visits ordered by season (left to right; Spring, Summer, Autumn, Winter).

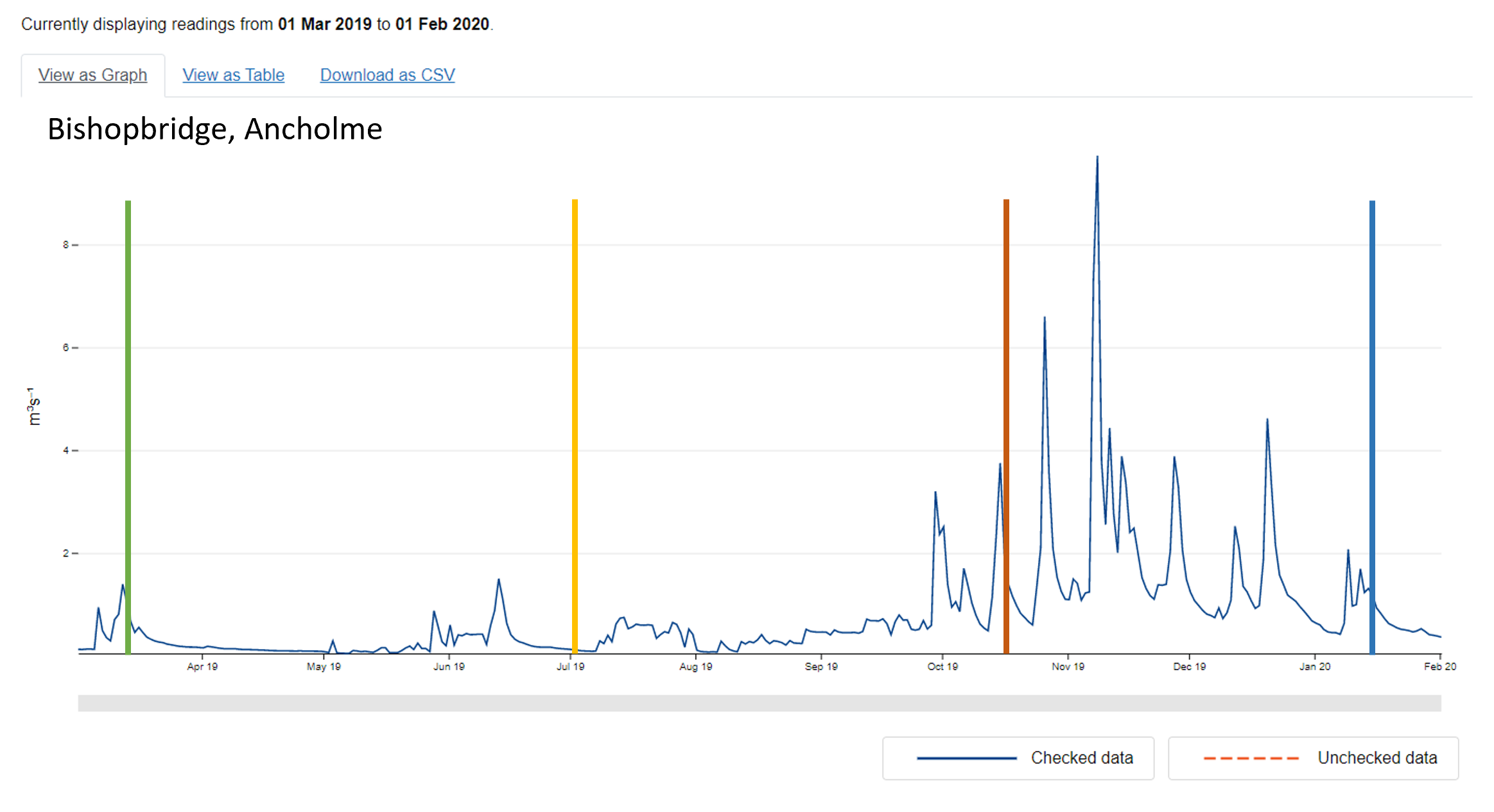

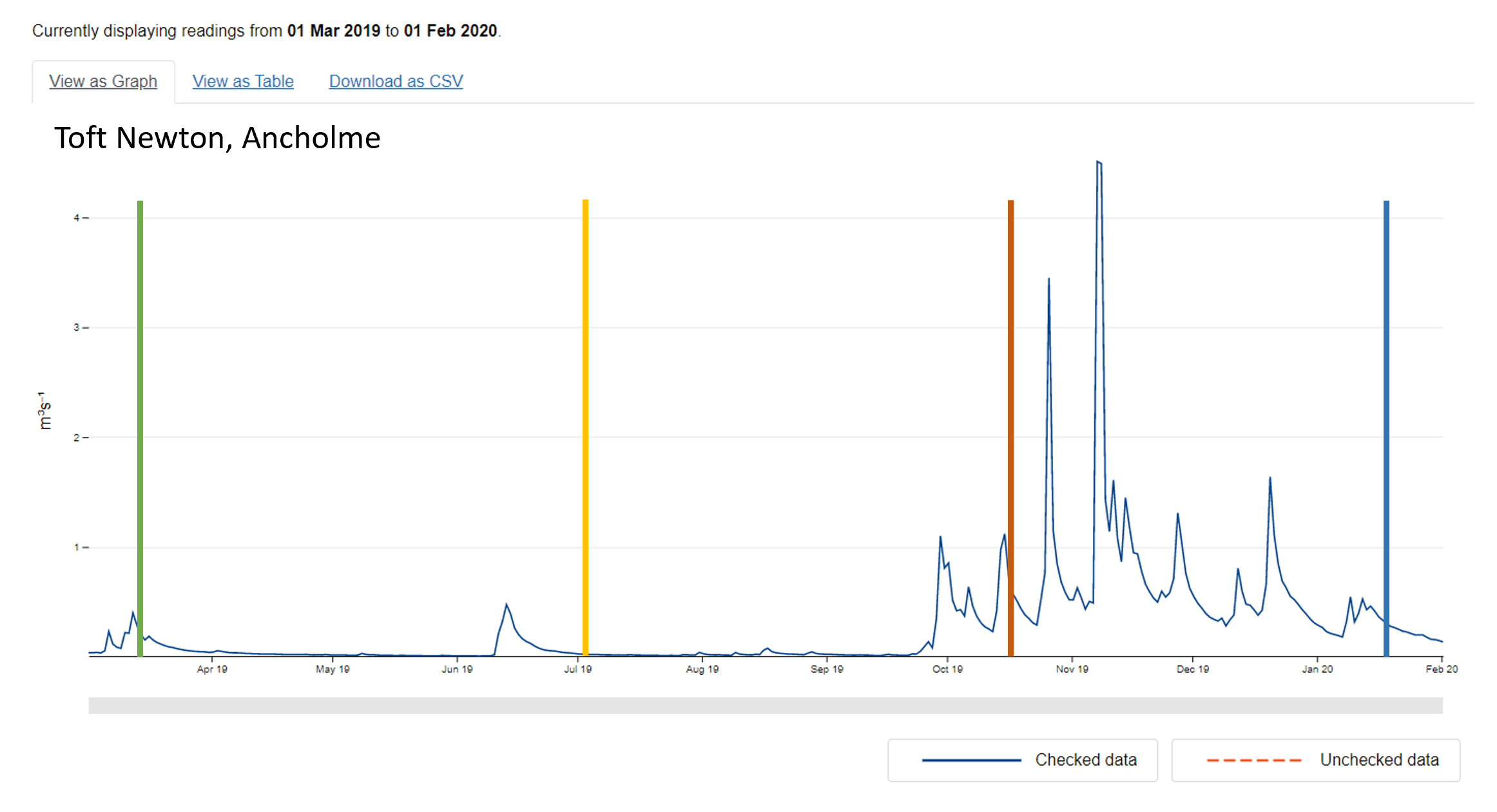

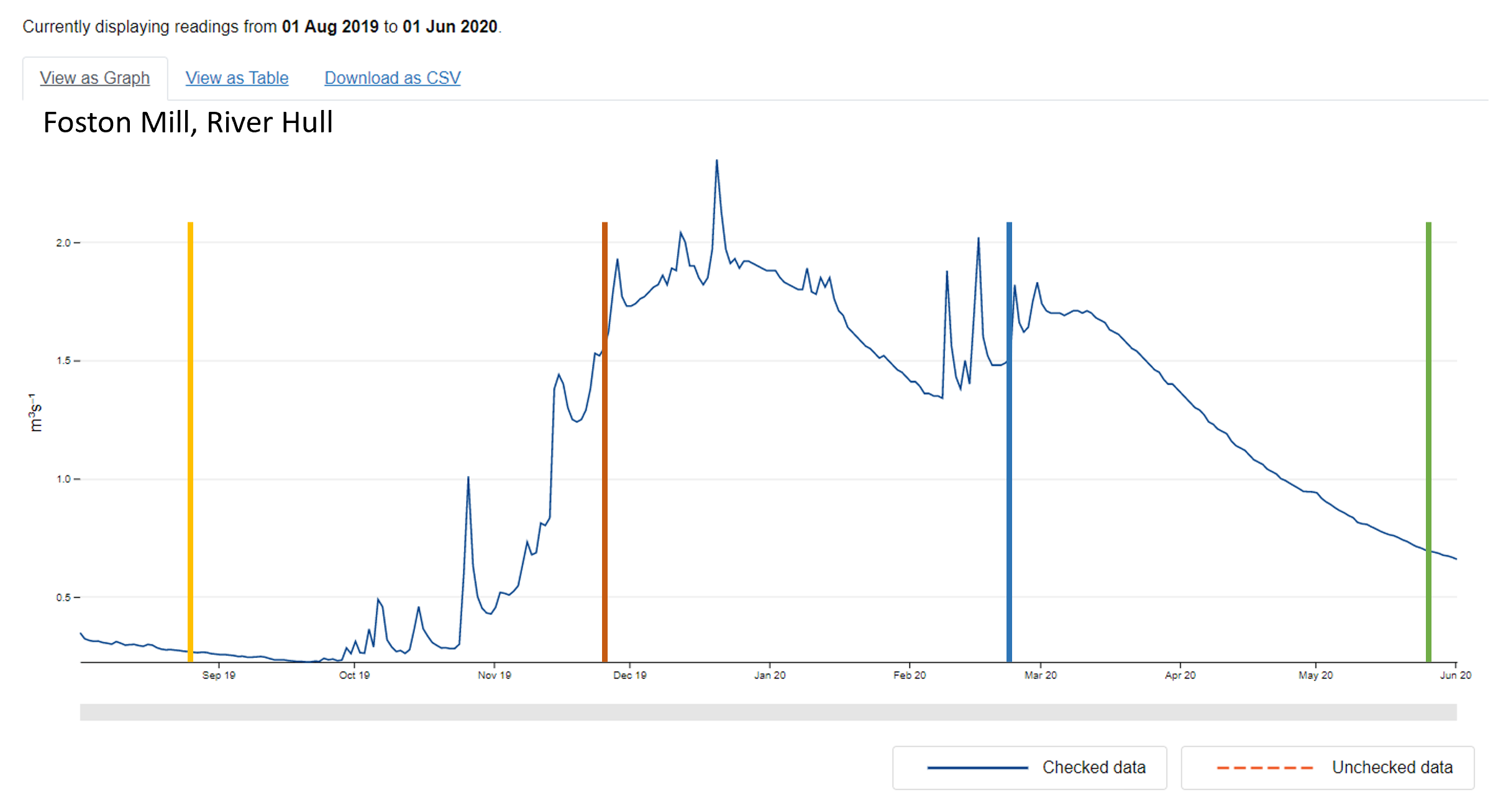

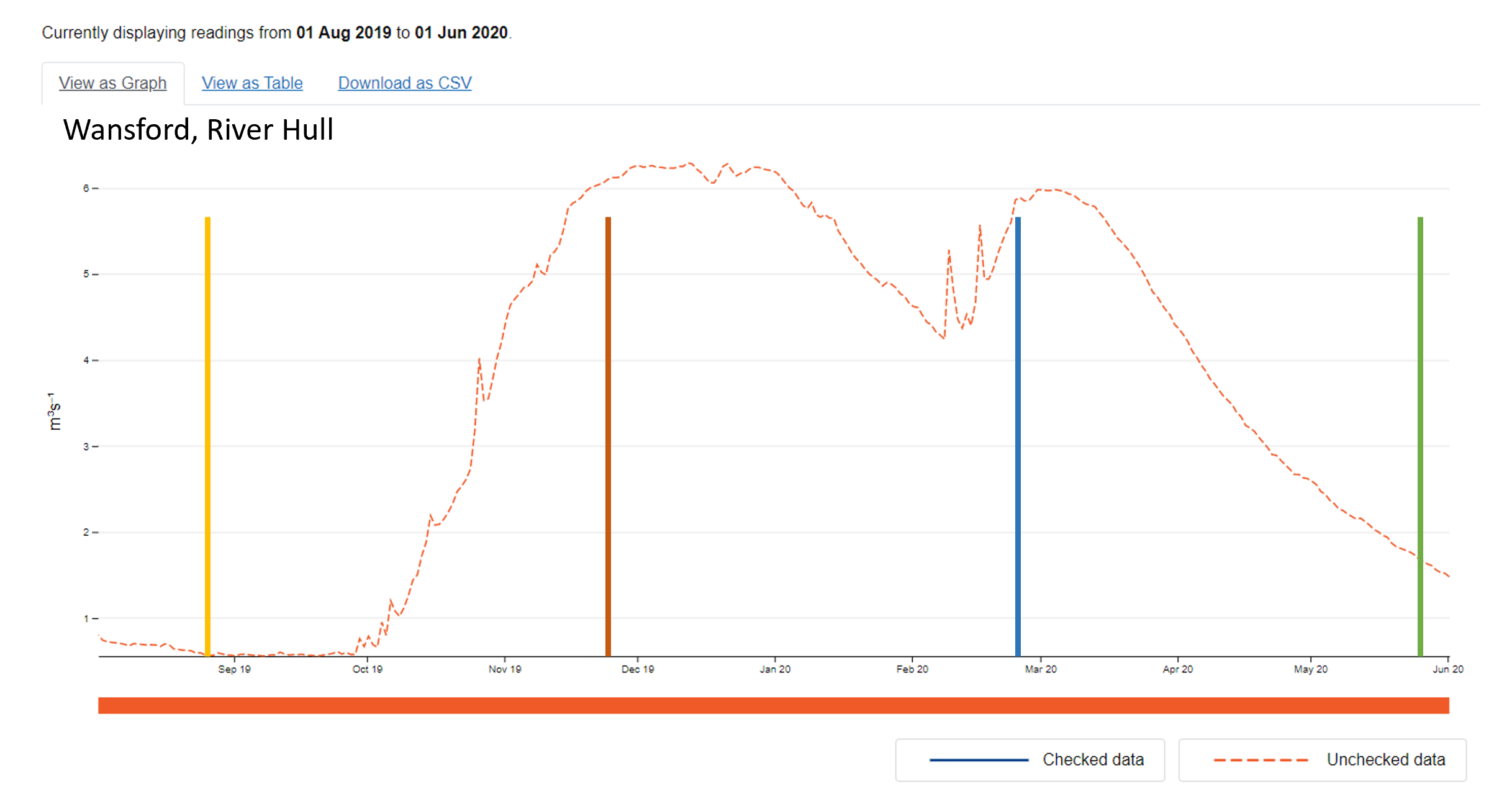
